# Modeling Mountain Pine Beetle Abundance and Distribution in a Changing Climate

**DOI:** 10.1101/2024.06.28.601294

**Authors:** Xiaoqi Xie, Micah Brush, Mark A. Lewis

## Abstract

The range of mountain pine beetle (*Dendroctonus ponderosae* Hopkins) is primarily constrained by climate, with winter temperatures playing a crucial role. Climate change is likely to increase the number of warm days and decrease cold days compared to historical norms, making higher latitudes more suitable habitats. In this work, we explore the potential impacts of environmental covariates on outbreaks of mountain pine beetle under climate change in a selected lodgepole pine area in Alberta. We employ a hierarchical model to examine mountain pine beetle dynamics approaching the end of the century. Our analysis assesses the impact of various climatic covariates and estimates the probability and expected number of infestations across different climate change scenarios. Our results from the hierarchical model underscore the critical role of degree days and overwinter survival probability, displaying an overall trend towards a higher probability and a greater number of outbreaks with increasing temperature. Our results indicate that Alberta is likely to experience widespread infestations in the future.

## 1 Introduction

The mountain pine beetle (MPB), *Dendroctonus ponderosae* Hopkins, is widely recognized as the most destructive bark beetle in North America, having killed more than 50% of the merchantable pines in British Columbia during the 2000s (Corbett et al., 2015). Although lodgepole pine is its primary host, MPB inhabits most species of pines within its habitats, where its range is mainly governed by temperature and seasonal suitability (Amman, 1978; Jenkins et al., 2001; Logan & Bentz, 1999; Logan & Powell, 2001; Safranyik & Carroll, 2007; Taylor & Safranyik, 2003). Climate change is facilitating MPB range expansion towards the east and north (Cullingham et al., 2011). The outbreak in the late 1990s and early 2000s enabled MPB to breach the Rocky Mountains into Alberta, thereby posing a threat to the boreal forest in Saskatchewan and beyond (Nealis & Peter, 2008; Robertson et al., 2009a; Safranyik et al., 2010).

Warm and appropriate temperatures across different seasons are important for MPB (Bentz et al., 1991). It usually completes its life cycle in one year, progressing through four life stages: egg, larva, pupa, and adult (Safranyik & Wilson, 2006). In the summer, MPB lays eggs that hatch into larvae. These larvae spend the winter and spring beneath the bark, emerging as adults the following summer. The adults then leave their hosts to find new ones. Once beetles successfully infest a new host tree, they cut-off the tree’s water supply system, causing it to turn red the following year. We call these trees that were killed last year and turn red this year ‘red-top’ trees. For a one-year life cycle, beetles typically require a cumulative temperature of at least 833 degree days above 5.6°C (Safranyik & Wilson, 2006). Most beetles emerge when temperatures exceed 20°C, and a range of 19°C to 41°C is necessary for successful dispersal in summer (Mccambridge, 1971; Safranyik & Carroll, 2007). Additionally, overwintering success is crucial for maintaining MPB populations (Aukema et al., 2008; Safranyik, 1978). The larval stage of MPB, typically occurring in winter, is recognized as the stage with the most cold tolerance (Bentz & Mullins, 1999; Safranyik & Wilson, 2006). Other life stages have lower cold tolerance, with the egg and pupal stages reaching 50% mortality at around *−*20°C, and adult beetles rarely survive the winter (Bleiker & Smith, 2019; Bleiker et al., 2017).

Climate change can transform previously unsuitable areas into good habitats, serving as a key driver behind the invasion of many species (Iverson & Prasad, 1998). Over the past three decades (1980-2010), the temperature has reached a historical high, with carbon dioxide (CO_2_) levels 40% higher than those in 1750 (Hartmann et al., 2013). Alberta has experienced a rise in summer temperatures ranging from +0.1 to +0.3°C per decade, accompanied by a significant increase in the number of days with temperatures above 25°C and a decrease in the frequency of cold days (Hayhoe & Stoner, 2019). Moreover, other studies shows that seasonal temperatures in Alberta are expected to increase by approximately 0.048°C annually, with the annual mean minimum temperature projected to rise from around -5°C to 0°C by 2100 (Eum et al., 2023; Jiang et al., 2017). Rising temperatures across all seasons lessen the challenge of accumulating the minimum degree days required for the beetle’s life stage development (Jiang et al., 2017). An increase in the number of days with temperatures above 25°C during summer leads to heightened beetle activity, facilitating greater dispersal (Safranyik & Wilson, 2006). The decrease in the frequency of cold days during winter significantly increases the overwinter survival rate for all beetle life stages, resulting in more beetles emerging in the subsequent summer (Régnière & Bentz, 2007). Rising temperatures also have the potential to directly increase forest productivity by providing extra heat and lengthening the growing season. However, this benefit could be negated if the increase in warmth is not accompanied by a sufficient rise in precipitation, potentially exacerbating drought conditions and adversely affecting tree growth and condition (Chhin et al., 2008; Monserud et al., 2008; Sauchyn et al., 2003).

Precipitation also stands out as a critical factor influencing the population dynamics of beetles. While seasonal precipitation could change from -25 to 36% in Alberta (Hayhoe & Stoner, 2019), the dryness index is expected to rise by 20% to 30%, suggesting a more pronounced drying trend (Barrow & Yu, 2005). Seasonal variations in precipitation can introduce further uncertainties regarding tree health, and the increased dryness could reduce pine trees’ resistance to beetle attacks.

Our objective was to study MPB dynamics in the context of global warming by creating risk maps. We generated these maps using a hierarchical model that includes both a presence model and an abundance model. Our research investigated the effects of various climatic variables on MPB dynamics under different climate change scenarios. Considering the expected changes in temperature and precipitation, we anticipate an increase in the frequency and severity of infestations within lodgepole pine forests.

## 2 Method

### 2.1 Study area and data preparation

To study the dynamics of MPB under climate change, we used infestation data from a square region with a side length of 50,000 meters in western Alberta, as depicted in Figure 1a, where the main MPB host trees are lodgepole pines. Our infestation data comprises data from aerial surveys, ground surveys, and control programs (Agriculture and Forestry, 2020; Sustainable Resource Development, 2007). In Alberta, these surveys are conducted to exhaustively search for infested trees within designated areas. Aerial surveys are carried out by observers in rotary wing aircraft counting red-top trees with Global Positioning Systems (GPS) to record locations. We refer to data from aerial surveys as ‘heli-GPS’ data. Ground surveys and control programs involve visiting the identified sites directly. Since ground surveys and control programs follow aerial surveys, we determined the grid cell size based on the distance between helicopter’s flight lines. Considering that the distance between two flight lines is approximately 500-1000 meters, we made the grid cell size to be 500 meters. An example of flight lines is depicted in Figure 1b. This area contains 10,000 cells, although the exact number of cells contained in each year’s data varies due to slight differences in each year’s detected area. Our analysis covered data from 2007 to 2020, a period that exhibits an increasing and then decreasing MPB population, as shown in Figure 1c. Previous research has found that the effects of climate or other covariates may change depending on the phase of population dynamics being studied (Kunegel-Lion & Lewis, 2020). Hence, we separated 14 years data into two stages, where the outbreak stage contains data from 2007 to 2014 and the growth peaking declining stage contains data from 2015 to 2020. Because our focus of research is future outbreaks of MPB, we focused our analysis in this paper on the outbreak stage. Details of the declining stage can be found in Xie, 2024.

**Figure 1:**
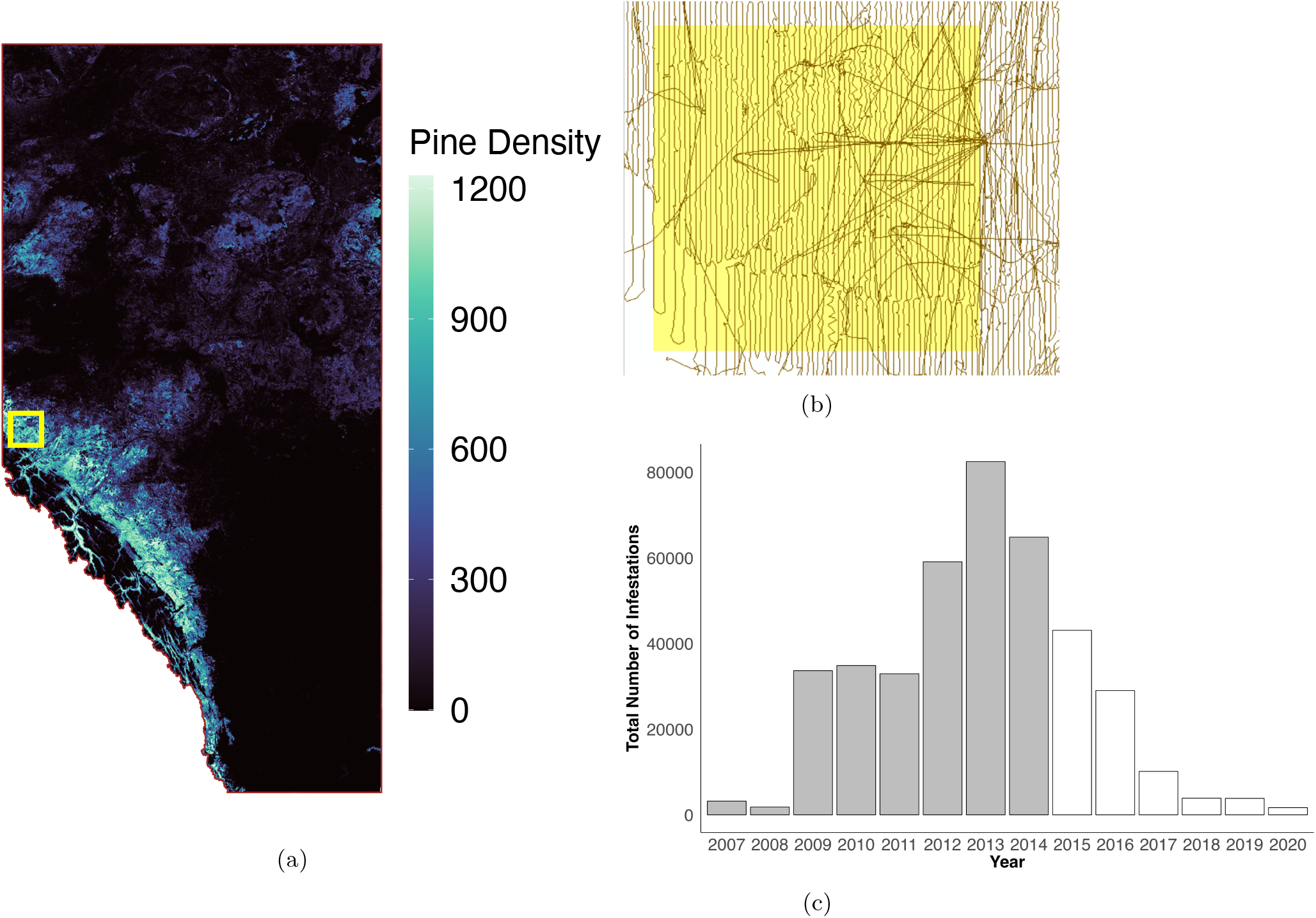
(a) Study area selected for analyzing the impacts of climate change, highlighted in yellow. Pine density is measured in stems per hectare, (b) 2019 helicopter’s flight lines represented by brown lines, with the selected area highlighted in yellow, and (c) Histogram depicting the observed total infestations within the selected area over a 14-year period, split into two stages. The outbreak stage is shown in grey (2007-2014) and the growth peak declining stage is shown in transparent (2015-2020).

As shown in Table 1, we considered climatic and ecological covariates, where each included covariate has a plausible biological mechanism for affecting pine beetle infestation. Details regarding covariates and their selection are given in Appendix A. Pine density indicates the number of host trees available for MPB and age reflect the vigor of lodgepole pines, indicating their resistance to MPB. Relative humidity accounts for the moisture content in the air, impacting both beetle and tree physiology. An increase in relative humidity can benefit trees, while its impact on beetles is uncertain. Wind speed provides insight into dispersal distances. Stronger winds can carry beetles to farther places. Soil moisture index reflects the land’s moisture content, which can influence tree health and, consequently, resistance to infestation (Hogg et al., 2013). Degree days affect the beetle’s brood development. MPB requires at least 833 degree days above 5.6°C to complete a oneyear life cycle (Safranyik & Wilson, 2006). Overwinter survival probability indicates larvae survival through the winter, estimated by a parameterized model as the chance of successful transition from the onset to the end day of winter (Régnière & Bentz, 2007).

**Table 1:**
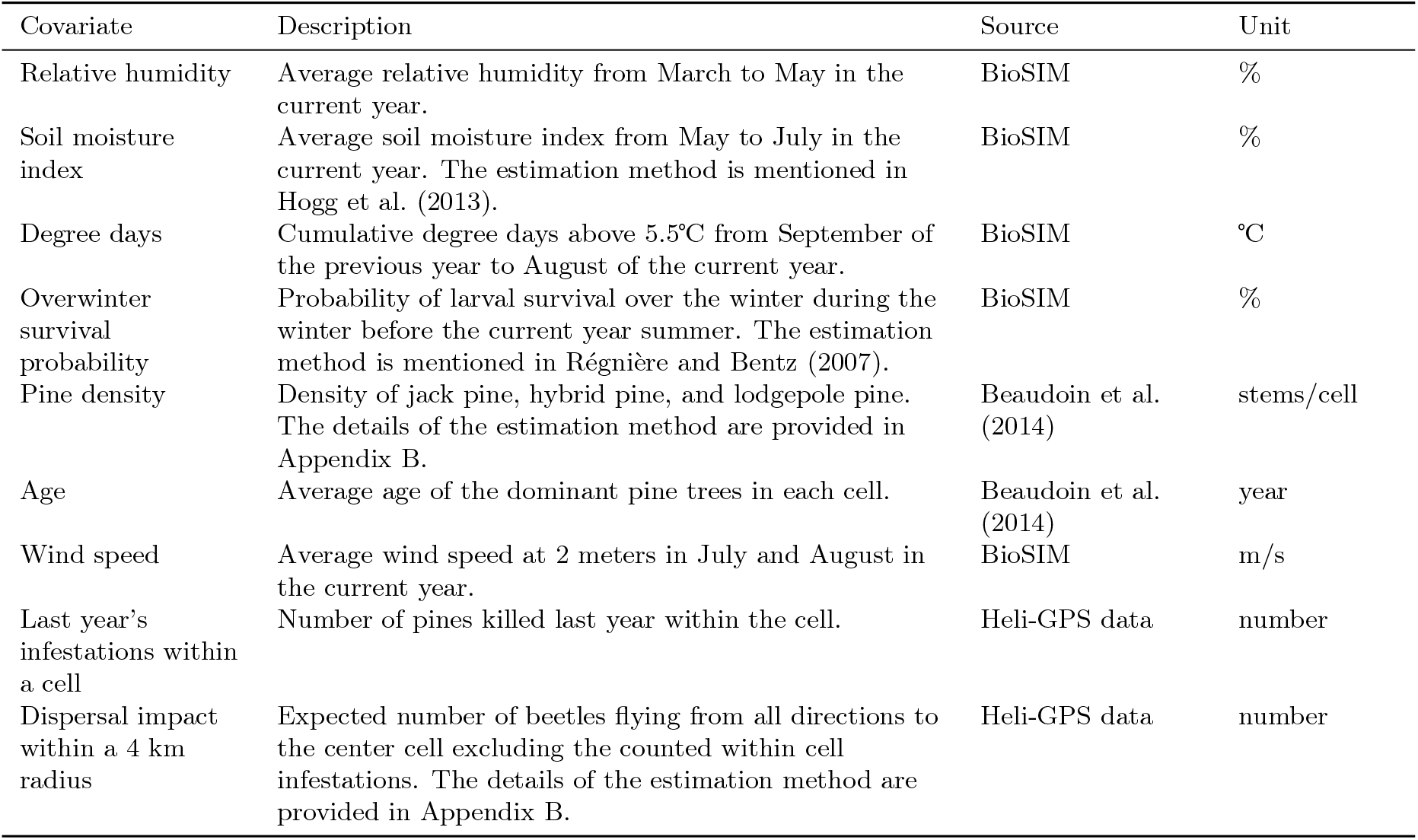
Overview of model covariates, with the first four rows highlighting four climatic covariates affected by global warming.

| Covariate | Description | Source | Unit |
| --- | --- | --- | --- |
| Relative humidity | Average relative humidity from March to May in the current year. | BioSIM | % |
| Soil moisture index | Average soil moisture index from May to July in the current year. The estimation method is mentioned in Hogg et al. (2013). | BioSIM | % |
| Degree days | Cumulative degree days above 5.5°C from September of the previous year to August of the current year. | BioSIM | °C |
| Overwinter survival probability | Probability of larval survival over the winter during the winter before the current year summer. The estimation method is mentioned in Régnière and Bentz (2007). | BioSIM | % |
| Pine density | Density of jack pine, hybrid pine, and lodgepole pine. The details of the estimation method are provided in Appendix B. | Beaudoin et al. (2014) | stems/cell |
| Age | Average age of the dominant pine trees in each cell. | Beaudoin et al. (2014) | year |
| Wind speed | Average wind speed at 2 meters in July and August in the current year. | BioSIM | m/s |
| Last year's infestations within a cell | Number of pines killed last year within the cell. | Heli-GPS data | number |
| Dispersal impact within a 4 km radius | Expected number of beetles flying from all directions to the center cell excluding the counted within cell infestations. The details of the estimation method are provided in Appendix B. | Heli-GPS data | number |

The last two covariates, last year’s infestations within a cell and dispersal impacts within a 4 km radius, imply the beetle pressure on nearby healthy lodgepole pines. The study conducted by Kunegel-Lion and Lewis (2020) demonstrates that the influence of an infested tree can be significant within a 1.25 km radius, while another report by Carroll et al. (2017) suggests that the potential impact of an infested tree over a 4 km radius could be less important. Therefore, in this analysis, we account for the influence of infestations within a 4 km radius, measuring the distance from the center of cells. We counted the exact number of infested trees within each cell and the expected number of beetles emerged within the cell from last year’s infestations in cells within a 4 km radius while excluding the center cell. The details about covariate derivation are presented in Appendix B.

The historical values of all climatic covariates, including relative humidity, wind speed, soil moisture index, degree days, and overwinter survival probability, were simulated in BioSIM using data from the four nearest weather stations. All other covariates were from existing maps, as described in Appendicies A and B. Additionally, we did not include some of the summer temperature covariates, such as maximum and minimum temperatures, because their effects on raised or lowered temperature could be captured well by the degree days covariate.

We assumed that the influence of climate change on relative humidity, soil moisture index, degree days, and overwinter survival probability is important for MPB dynamics. The influence of climate change on other covariates were assumed to be less important. Their predicted value under global warming in 2100 were generated in BioSIM using the predicted weather data from the HadGEM2 model with a 10 km resolution (Martin et al., 2006; Ringer et al., 2006). We used three climate change scenarios: Representative Concentration Pathway (RCP) 2.6, 4.5, and 8.5. A higher RCP level indicates a scenario of more severe global warming, reflecting greater greenhouse gas emissions. In BioSIM, we produced 10 replications for each affected climatic covariate under each RCP scenario and then used the mean of these 10 replicates to replace the original covariates. These projections are presented in Figure 2. Under all three RCP scenarios, relative humidity was higher than the average value observed in the outbreak from 2007 to 2014. Higher RCP values led to higher relative humidities. Conversely, the trends for soil moisture index are opposite. The historical mean values are higher than those projected under climate change scenarios. With a rising future temperature, the soil moisture index decreases. Degree days exhibit an increasing trend under global warming. Historical average value are the lowest and as temperatures rise, degree days increase. The overwinter survival probability also follows an increasing trend. A higher overwinter survival probability is expected with a higher temperature.

**Figure 2:**
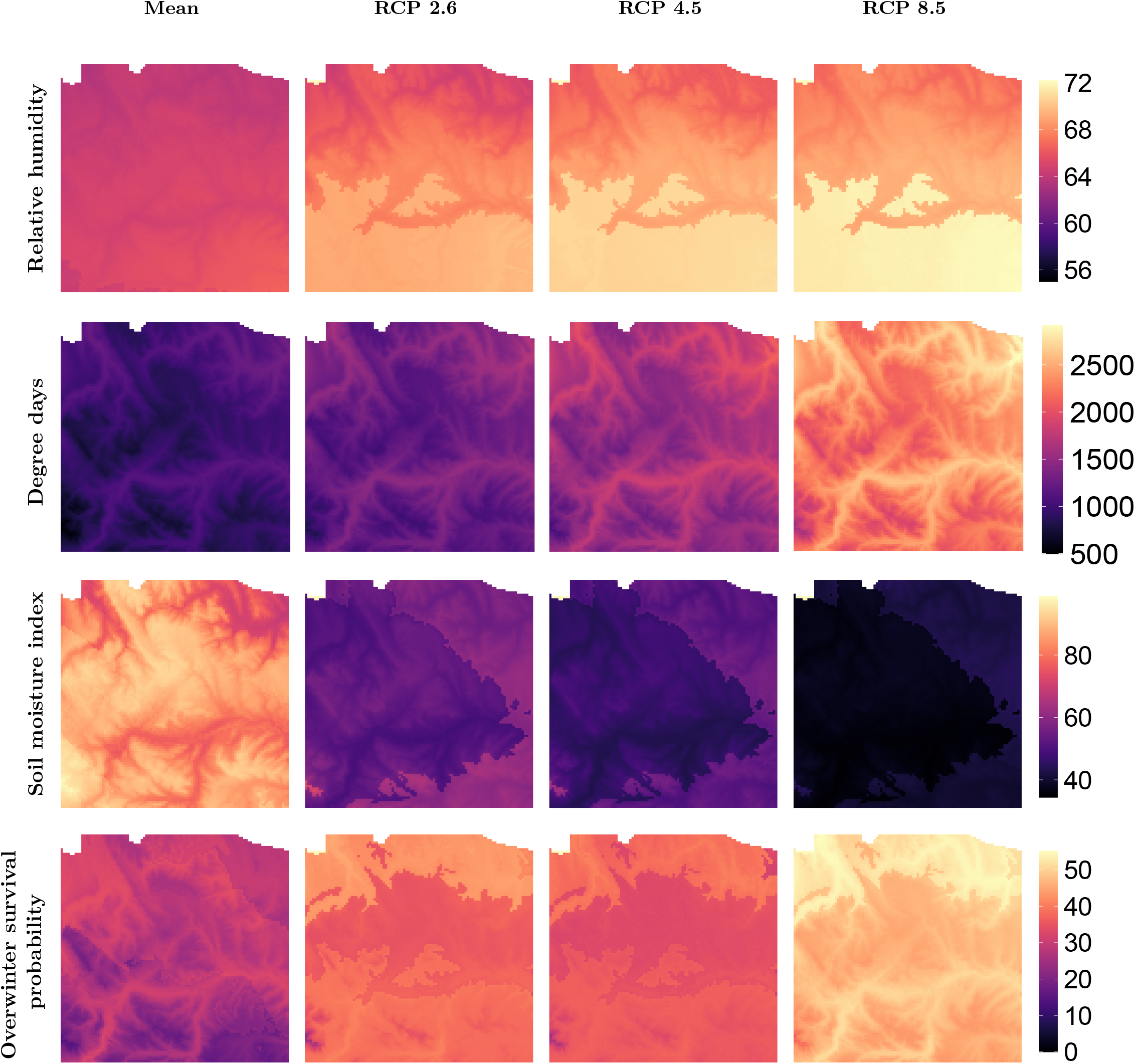
Comparisons of the trends of four climatic covariates—relative humidity, soil moisture index, degree days, and overwinter survival probability—across four scenarios, including the mean value of the outbreak stage (2007-2014) and three RCP levels (RCP 2.6, 4.5, and 8.5) in 2100. A cell value is estimated by averaging this cell values over years in the outbreak stage and 10 replications.

### 2.2 Statistical model

In this analysis, we examined the impact of four climatic covariates—relative humidity, soil moisture index, degree days, and overwinter survival probability—on MPB dynamics under climate change scenarios. Utilizing data from a dynamic stage, we applied the hierarchical Zero-Inflated Negative Binomial (ZINB) model. The hierarchical model comprises a presence model employing a Bernoulli distribution and an abundance model employing a Negative Binomial distribution. The presence model estimates the probability of that the beetles could be present, while the abundance model calculates the expected number of infestations, given that the beetles could be present. Note that zero infestations could arise either from a zero in the Bernoulli presence model or from a zero in the number arising from the Negative Binomial abundance model.

We use *Y* to represent the response variable showing the number of trees infested in the current year, and we use *Z* and *X* denote the sets of spatially indexed environmental covariate matrices that provide input for the presence and abundance models, respectively. In our case, we initially considered similar covariates for two models, meaning that the sets *Z* and *X* were equal. We designate *γ* and *β* to be the coefficient vectors for the presence and abundance models, respectively. We use the quantity *π* to signify the event of no infestation for the presence model, as determined by a Bernoulli distribution. The function *f*_Bernoulli_ denotes the probability density function for this Bernoulli distribution. We use the quantities *μ* and *ϕ* to represent the mean number of infestations and the dispersion parameter in the abundance model, as determined by the Negative Binomial distribution. The hierarchical ZINB model is then written as:

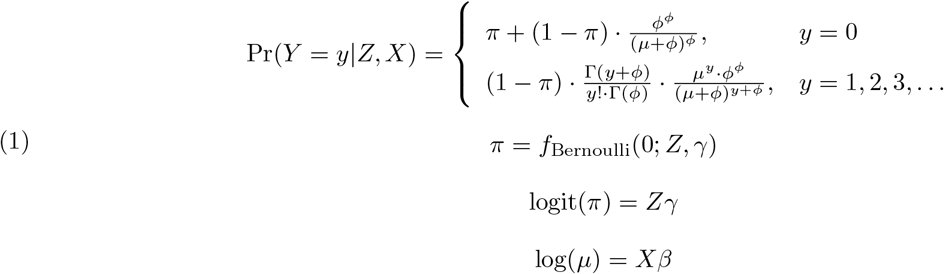

The model was fit to the spatial oubreak data (from 2007 to 2014) using Ridge Regression to address potential multicolinarity of environmental covariates, *Z* and *X*, and a lasso-type penalty given by Smoothly Clipped Absolute Deviation (SCAD) to address selection of variables (Appendix A).

We initially considered models nested within the ZINB, including the Poisson and Zero-Inflated Poisson models, but determined that the ZINB was the most effective (Appendices C and D). The effectiveness of each model was determined through model selection (BIC, Appendix C), model fit (Randomized Quantile Residual, Appendix C) and model validation (Appendix D). Model validation assessed the performance of one and two year predictions via accuracy and root-mean-square error (Appendix D).

### 2.3 Comparative analysis under climate change

We assessed the risk by projecting the probability of infestations using the presence model and the expected number of infestations through the abundance model for the year 2100 under each RCP scenario. We examined three RCP scenarios: 2.6, 4.5, and 8.5. To focus solely on the potential effects of four climatic covariates on new infestations under climate change, we fixed all unaffected covariates except beetle pressures to be the median values for each cell to avoid the influence of extreme values. We aimed to maintain beetle pressures at a moderate level, setting the values for the last year’s infestations within a cell and the dispersal impact within a 4 km radius at the 10th percentile of the positive values in our dataset, which were 3 and 0.05, respectively. The four climatic covariates were replaced with the average value from 10 simulations generated in BioSIM using the predicted weather data from the HadGEM2 model (Martin et al., 2006; Ringer et al., 2006). Thus, during a dynamic stage, for each RCP scenario, we created five maps for both the presence and abundance models: four maps were generated by replacing one affected covariate at a time, and in the fifth map, all four covariates were replaced. Notably, when replacing just one covariate, we also set the other three covariates to the median values of each cell. For each cell, we have historical data and we took the median of these values during a dynamic stage. The general procedures are depicted in Figure 3.

**Figure 3:**
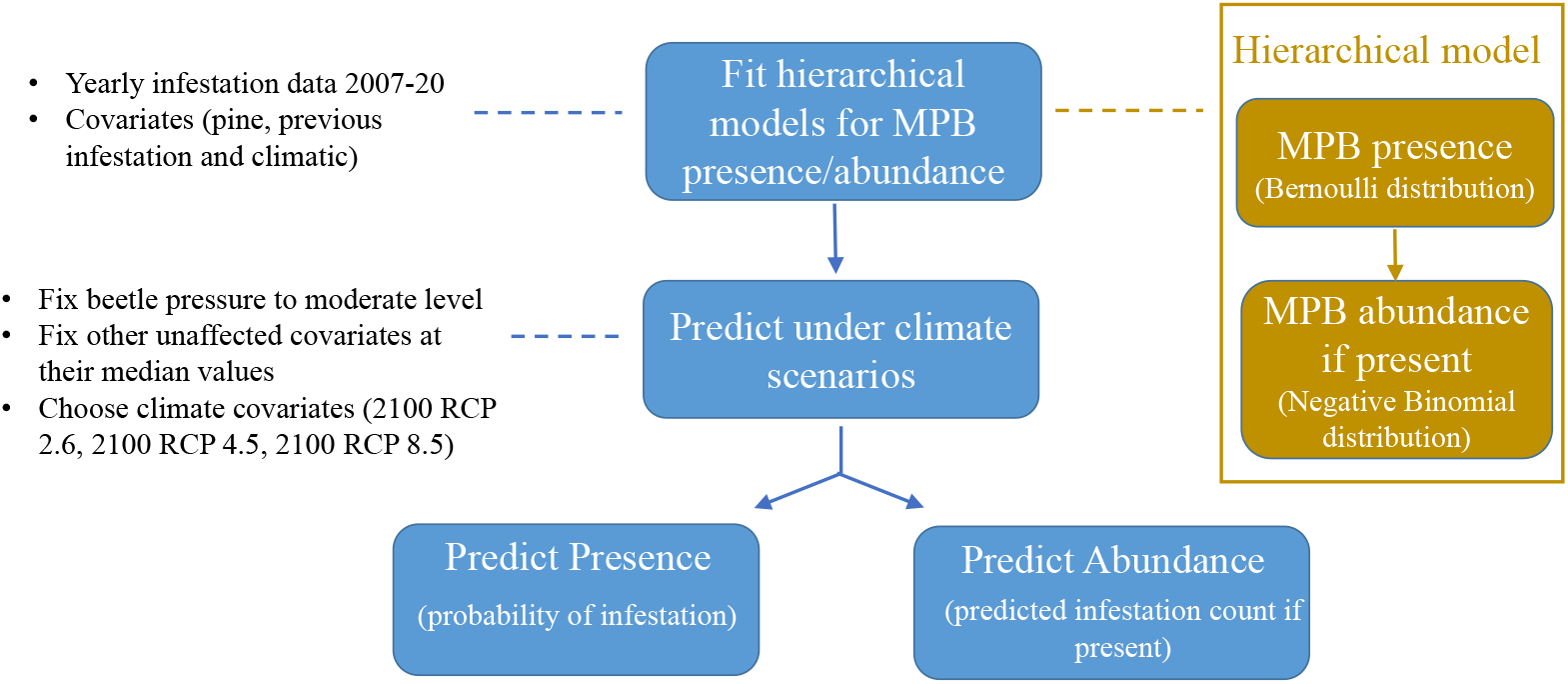
Analysis for assessing the impacts of four climatic covariates: relative humidity, soil moisture index, degree days, and overwinter survival probability. Beetle pressures were maintained at a moderate level, while the remaining covariates except four climatic covariates, presumed stable under climate change, were set to their median values.

## 3 Results

Temperature can have a positive impact on beetles, while the influence of moisture content can benefit both beetles and trees. The presence model corresponds to the probability of infestations, whereas the abundance model concerns the number of infestations. In the fitted hierarchical model, a positive coefficient suggests either a higher probability or a greater number of infestations. In this paper, our focus is on future outbreaks of MPB, so we only presented the results in the outbreak stage as illustrated in Figure 4. The rest of results in the growth peak declining stage can be found in Xie, 2024. For the four climatic covariates, the coefficients in both models demonstrated that overwinter survival probability and degree days have a positive effect on beetles, while relative humidity negatively affects beetles. Furthermore, overwinter survival probability and degree days were identified as the most significant among the four climatic covariates. The impacts of soil moisture index were ambiguous as its 95% confidence intervals include 0. For the other covariates, a higher value of last year’s infestations within a cell, wind speed, and dispersal impact within a 4km radius were found to benefit beetles, leading to a higher probability and number of infestations. Pine density positively impacted the presence model, indicating that the number of lodgepole pines within a cell is positively correlated with the probability of infestations. However, its impact in the abundance model on the number of infestations was unclear, as its confidence interval included 0. Age can negatively affect both the presence and abundance models. The covariate representing last year’s infestations within a cell was identified as the most influential covariate, underscoring the impact of nearby infestations on healthy lodgepole pines.

**Figure 4:**
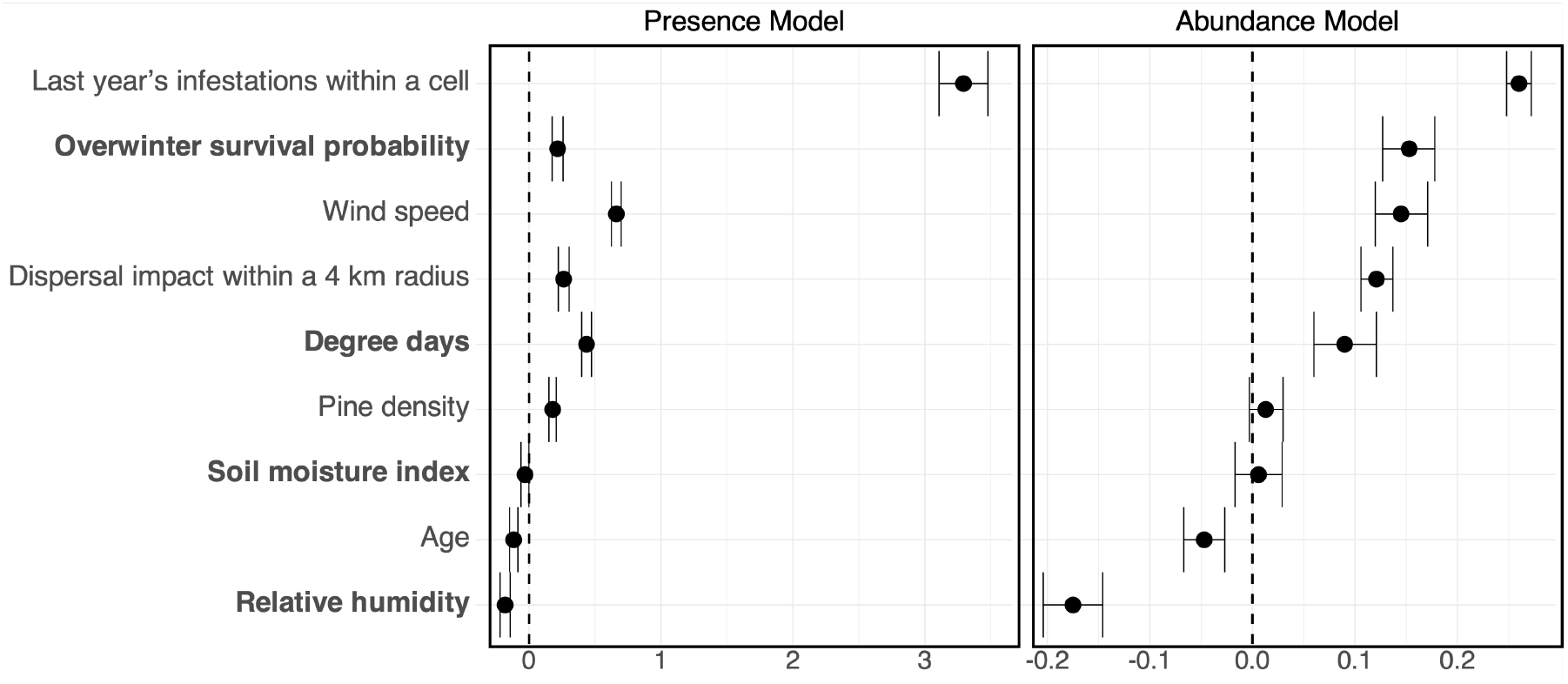
The first plot presents coefficients of standardized covariates from the presence model, and the second plot from the abundance model of the hierarchical model for the specified region (see **Figure 1a). These are shown during the outbreak stage with four climatic covariates affected by global warming in bold. Covariates are ordered by their coefficient values in the abundance model.

The projections under climate change indicated that global warming will likely benefit beetles leading to widespread infestations. It is important to note that the expected number of infestations is presented on a logarithmic scale. Our results showed that changes in relative humidity lead to a decrease in both the probability and number of infestations, as illustrated in Figures 5 and 6. The decrease from the current scenario to RCP 2.6 was notable, yet the differences amongst the three RCP scenarios were less obvious, especially between RCP 4.5 and 8.5. Figures 5 and 6 also demonstrated that an increase in degree days significantly raises both the likelihood and number of infestations. A clear increase in infestation probability was observed from the current scenario to RCP 2.6, and the result under RCP 8.5 showed that most cells achieve the highest probabilities when temperature reaches the highest expected level. For the number of infestations, the increase from the current scenario to RCP 2.6 was less pronounced than that for infestation probability. The result under RCP 8.5 still showed a significant increase compared to results under other scenarios. The influence of soil moisture index under climate change on beetles was limited. The plots showing the probability and number of infestations with respect to this feature did not present significant difference under different RCP levels. This is because the effect sizes, as given by the standardized coefficients are close to zero in our model. The impact on overwinter survival probability from climate change was relatively minor. Figures 5 and 6 revealed a slight decrease in both the likelihood and number of infestations when transitioning from RCP 2.6 to RCP 4.5. However, moving from RCP 4.5 to 8.5, we saw an increase in both values. By substituting all four climatic covariates simultaneously, we noted an increase in both the probability and number of infestations as the global mean temperature rises, as shown in Figures 5 and 6. The projected results in both models under RCP 2.6 were closest to current values. The results under RCP 8.5 showed the largest increase compared to the current values. Our findings suggested that under the warmest scenario, lodgepole pine forests in Alberta could face widespread infestations.

**Figure 5:**
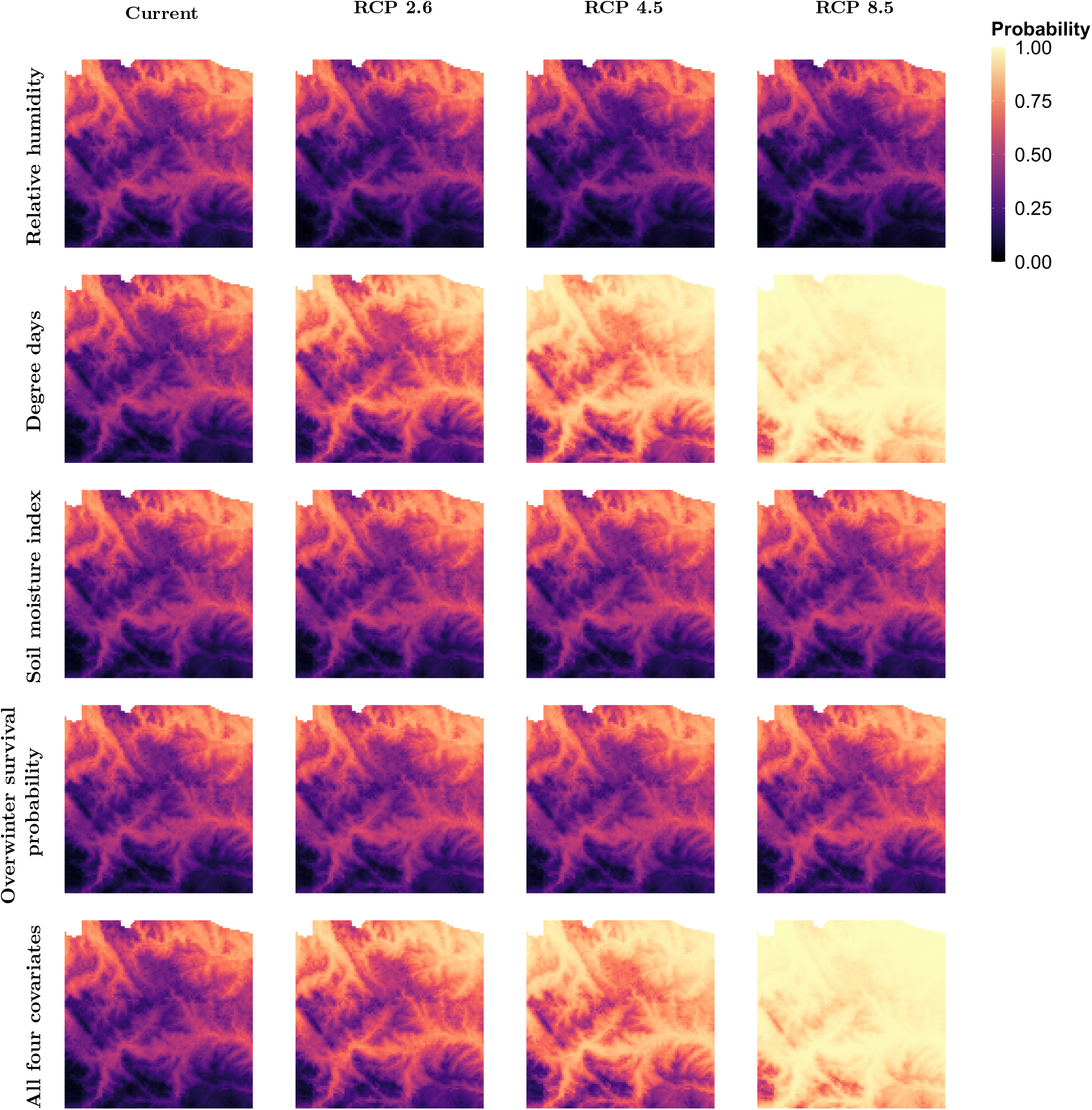
Comparisons of current and projected infestation probabilities in 2100 under three RCP levels, as estimated by the presence model. The current probability was calculated using median values of covariates, excluding last year’s infestations within a cell and dispersal impacts within a 4 km radius. They were fixed at a moderate level of beetle pressure. Last year’s infestations within a cell and dispersal impacts within a 4 km radius were fixed at 3 and 0.05. The current probabilities depicted across different rows corresponded to the same plot. In the first four rows, the value of one climatic covariate was replaced under global warming conditions at a time, whereas in the last row, all four climatic covariates were replaced.

**Figure 6:**
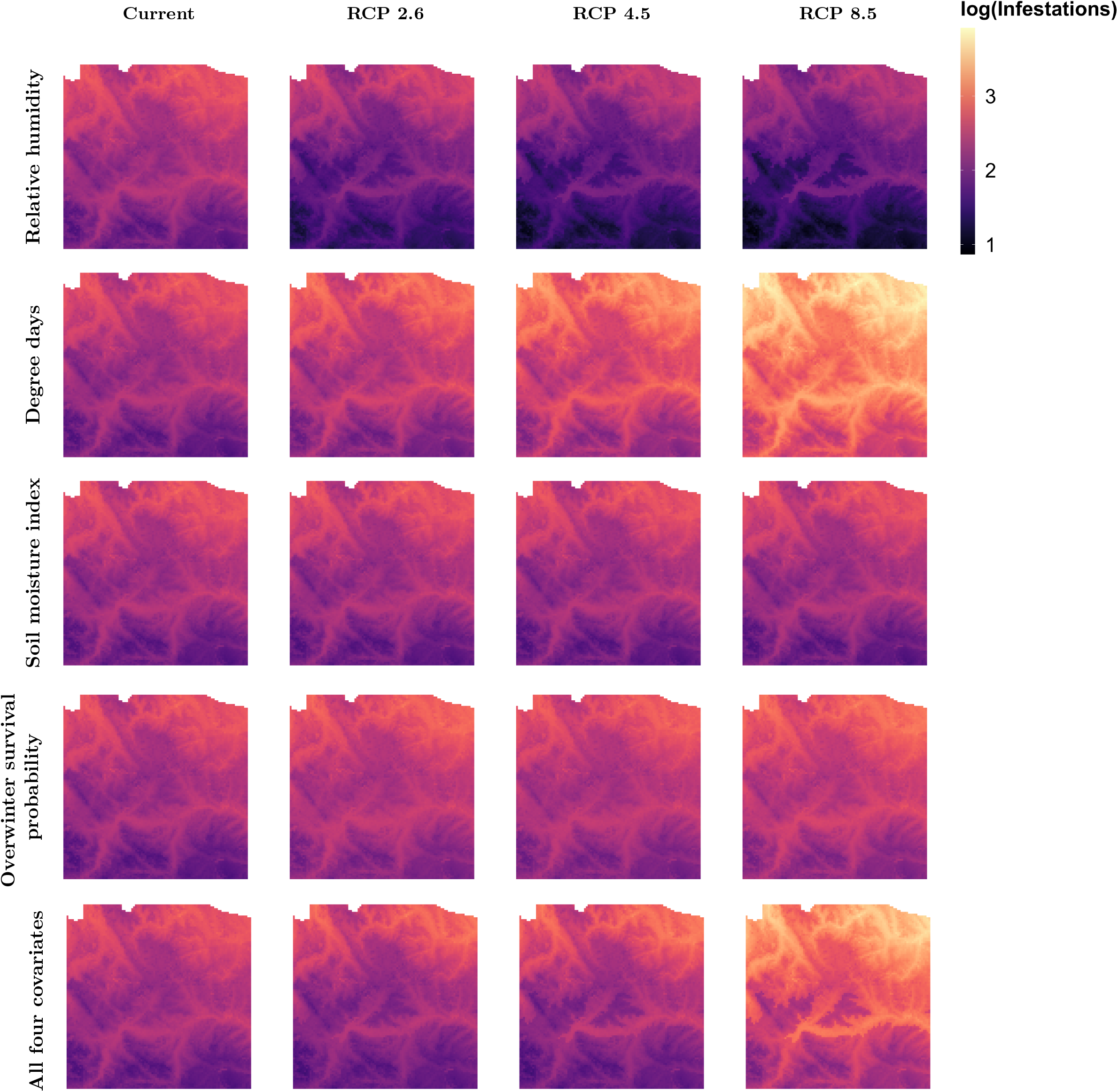
Comparisons of current and projected expected infestations in 2100 under three RCP levels, as estimated by the abundance model. It is important to note that the expected number of infestations is presented on a logarithmic scale. The current expected number was calculated using median values of covariates, excluding fixed last year’s infestations within a cell and dispersal impacts within a 4 km radius. Last year’s infestations within a cell and dispersal impacts within a 4 km radius were fixed at 3 and 0.05. The current expected number depicted across different rows corresponded to the same plot. In the first four rows, the value of one climatic covariate was replaced under global warming conditions at a time, whereas in the last row, all four climatic covariates were replaced.

## 4 Discussion

Our analysis revealed possible effects of climate change in the population dynamics of MPB. In our model, degree days emerged as the most critical factor. Its significant coefficients, combined with increased values under climate change, directed our projections for both the probability and number of infestations, as depicted in Figures 5 and 6. Its positive coefficients suggest that degree days help beetles, leading to a higher likelihood and number of infestations. This finding aligns with biological understanding that higher degree days benefit brood development and is an indication of a warmer year (Safranyik & Carroll, 2007). Overwinter survival probability, while also crucial, had a lesser impact on infestations compared to degree days. A higher overwinter survival probability is a consequence of a warmer winter. Its positive coefficients in the hierarchical model indicate that a milder winter is an important factor for the MPB population, which supports previous studies that identified winter temperature as a significant constraint on beetle population expansion (Aukema et al., 2008; Logan et al., 2003; Preisler et al., 2012; Robertson et al., 2009a; Sambaraju et al., 2012a). According to simulations from BioSIM, the average summer relative humidity showed an overall increasing trend, albeit with local variations. This rise in relative humidity, coupled with its negative impact on beetles, generally reduced both the probability and number of infestations. The projections suggest that increased relative humidity could aid lodgepole pines improving their resistance to beetle attacks. The impact of the soil moisture index remained ambiguous in our model, as its confidence intervals in both presence and abundance models included zero. Previous work has found that the effect of precipitation variables may depend on moisture in years previous to the year of infestation, or cumulative drought (Preisler et al., 2012), which may explain the small and ambiguous effect of these variables in our model. Outside of climate covariates, the small negative coefficients of age in both presence and abundance models indicate that as trees age, both the probability of infestations and the expected number of infestations decrease, implying stronger resistance to MPB. This aligns with findings that trees up to a certain age are likely more resistant to MPB infestation (Safranyik & Wilson, 2006). However, it is also likely that older trees with thicker phloem produce more beetles (Safranyik et al., 1975; Safranyik & Wilson, 2006), so we might expect the sign of the age coefficient to be positive in the abundance model.

Our results offered insights into population dynamics under global warming but should not be considered to be definitive predictions. While we validated over short time scales, we do not expect the model to have good predictive capacity over longer, decadal scales. Rather, the model provides projections of possible outcomes over longer time scales, based on the quantitative relationships determined between the climatically influenced environmental covariates and the presence and abundance of beetles, as parameterized over shorter time scales. There is also the issue of extrapolating our regression models to project population dynamics for environmental covariates that lie outside the training data range. While we expect that the overall trends, described in our model fits to data from observed from 2007 to 2014, to continue outside of this range, we have not shown this definitively. In other words, we are projecting how a climatically influenced environmental covariate will affect beetle presence and distribution under climate change based on similar, but not as extreme, situations as in the past.

Our model focuses on the influence of expected changes in relative humidity, soil moisture index, degree days and overwinter survival probability. However, climate change may potentially increase the variability of extreme weather events, potentially leading to significant beetle mortality within brief periods. Early cold snaps, late cold snaps, and sudden drops in temperature can adversely affect beetles, thereby reducing the likelihood of outbreaks (Sambaraju et al., 2012b). Extreme weather was not considered in our study as we only took means of climate projections.

With respect to the actual climate change predictions that are used in our modelling, the HadGEM2 model is known for its conservative projections. This suggests that the actual impact of climate change could be bigger than our projections indicate (Collins et al., 2011).

Finally, our findings are consistent with previous studies indicating an increased likelihood of outbreaks under global warming (Aukema et al., 2008; Sambaraju et al., 2012a; Srivastava & Carroll, 2023). While past studies has explored how climate change might alter MPB habitats and expand their range, our study examined the effects of key climatic covariates on MPB dynamics in lodgepole pine forests, offering more insights into how specific factors will affect MPB activity. A warmer year accompanied by a drier summer could favor beetles and harm trees, leading to more severe infestations. A warmer year with a wetter summer could introduce uncertainty as both beetles and trees take advantages. The possibility of extreme weather conditions, especially in winter, also add uncertainty. A warmer year with frequent extreme winter weathers could result in fewer beetles emerging the following summer. In previous studies, seasonal temperatures are expected to increase by approximately 0.048°C annually, with the annual mean minimum temperature projected to rise from around -5°C to 0°C by 2100 (Eum et al., 2023; Jiang et al., 2017). These increases in temperature will lead to higher degree days and overwinter survival probability. These projections, combined with an increase in the dryness index by 20% to 30%, suggest that Alberta will become a more favorable habitat for beetles (Barrow & Yu, 2005).

## Supporting information

This contains all appendices.

## Acknowledgements

We would like to thank all members of the Lewis Research Group, particularly Evan Johnson and Kévan Rastello, for their comments on this project. We would also like to thank the members of the TRIA-FoR project for their support. We thank Kelsey Gritter for her help with data wrangling. Special thanks go to the Forestry Division, Government of Alberta, especially David Strauss and Caroline Whitehouse, for their assistance with our data. XX acknowledges the use of ChatGPT for assistance in checking grammar and vocabulary.

## Author contributions

Conceptualization: Mark A. Lewis;

Methodology: Mark A. Lewis, Micah Brush, Xiaoqi Xie;

Formal analysis and investigation: Xiaoqi Xie;

visualization: Xiaoqi Xie;

Software: Xiaoqi xie;

Supervision: Mark A. Lewis;

Writing – original draft: Xiaoqi Xie;

Writing – review & editing: Micah Brush, Mark A. Lewis, Xiaoqi Xie;

## Data availability

Data are not made available due to data sharing agreement at this moment.

